# Pillbox: A Leakage-Aware Foundation-Model Predictor and Lineage-Ceiling Diagnostic for Cancer Drug Response

**DOI:** 10.64898/2026.06.08.725572

**Authors:** Justin Hill, Eric Jiao, Shwetank Singh, Arhant Ghanta, Declan Anders, Joo Ho Jeong, Hong Joo Ryoo

## Abstract

We present Pillbox, a predictor whose pipeline is audited against the six Asiaee leakage modes with the one residual pathway shown by per-fold ablation to be non-load-bearing on hard splits. Our model combines CpGPT methylation embeddings, CLAMP drug embeddings, and per-fold-fit gene-expression principal components which are fused by Feature-wise Linear Modulation (FiLM)-conditioned graph attention on the STRING v12 protein-protein interaction graph. Then we *α*-ensemble the model against a histogram-based gradient boosting regressor baseline. On GDSC GSE68379 (987 cell lines, 375 drugs) across seeds 42, 7, and 123, the ensemble reaches test R^2^ of 0.78, 0.77, and 0.76 on random, histology-blind, and site-blind splits respectively, with cell-aware lifts above the drug-mean floor of +0.054, +0.060, and +0.037. As a quantitative diagnostic for feature-stack saturation we propose the cross-architecture residual correlation, calibrated against a same-architecture-different-initialization control. On histology-blind splits the cross-architecture value of 0.939 falls short of the same-architecture ceiling of 0.974 by approximately 0.03 in residual correlation, a gap we interpret as the headroom available to architecture choice on top of the current foundation-model representation and consistent with the long-established observation that tissue lineage dominates cell-line drug response. We integrated curated mutation, methylation, and drug-target-expression channels, but these do not improve prediction once foundation-model embeddings are in place. Cross-screen validation against PRISM matches the GDSC-to-PRISM measurement reproducibility ceiling within 0.01 Spearman.

## 1 Introduction

Predicting how a cancer cell line will respond to a drug from its molecular profile is one of the oldest open problems in pharmacogenomics. The clinical motivation for a machine learning driven approach is that a model trained on a panel of well-characterized lines should help nominate compounds for cancer types it has not been trained on. However, the reality is that published methods as of current have accumulated an evaluation problem which, by the most recent audit, affects the majority of them [1].

Three audits in the last eighteen months have made the situation clearer. Asiaee and colleagues [1] examined 32 published methods cited more than 3,000 times in aggregate, found data leakage in 23, and showed that leakage-free cross-validation raises mean squared error by 16.6 percent on average. The selected feature sets differ between leaky and corrected pipelines at Jaccard 0.18, which means the biomarkers reported in those papers are largely artifacts of the preprocessing protocol. Codicè et al. [2] introduced a sharper observation. A predictor that simply memorizes each drug’s training-set mean ln(IC_50_) reaches pair-pooled Pearson correlation of 0.85 on warm-start GDSC random splits. We replicate this exactly: a DummyDrugAvg baseline on our evaluation reaches Pearson 0.852 on random, 0.850 on histology-blind, and 0.844 on site-blind. The published 0.94 numbers are barely 0.09 Pearson above the drug-mean floor. Garai and colleagues [3] benchmarked five methods against a mean-based baseline and concluded that several of them cannot reliably outperform the baseline; Chen and Zhang [4] showed that under cold-drug evaluation a similarity-regularized matrix factorization baseline outperforms every deep learning method tested.

Standardized benchmarking efforts [5, 6] have begun to set a more honest floor. They are useful, and we follow the leakage-discipline principles they prescribe. However, neither effort ships a new prediction method that pairs foundation-model embeddings with that discipline. Our paper seeks to fill this gap.

The contribution of the work is as follows. We first provide a leakage-aware cancer drug response predictor with explicit verification against the six leakage modes catalogued by Asiaee. The residual M2 leakage pathway flagged by our internal audit (the global-fit gene-expression PCA plus the similarity-augmentation *k*-NN graph, six `fit()` calls total) is shown to be non-load-bearing on hard splits via a per-fold selective-removal / ablation. Second, we fill a specific empty cell in the post-audit competitive landscape: of the six methods Asiaee certified clean as of May 2026 (Velodrome, UNO, DeepResponse, ScreenDL [7], DrVAE, and MMDRP [8]; see Supplementary Table S6 for the full roster with citations), none combines a methylation foundation model with a drug foundation model under tissue blind hard splits and multi-seed validation. Third, we propose cross-architecture residual correlation as a quantitative diagnostic for feature-stack saturation. The lineage-dominance observation has been around since the original GDSC analysis [9] and has been re-articulated in recent work [10, 4], but a number for the ceiling it imposes has been missing, and we provide one. Fourth, we report a negative result: curated mutation, methylation, and drug-target-expression channels do not improve prediction once foundation-model embeddings are in place, the simplest explanation being that the embeddings already encode the lineage axis these priors carry.

The structure of the paper follows. Section 2 introduces the dataset, evaluation protocols, and modeling framework. Section 3 presents the main results and a series of analyses probing robustness, interpretability, and architectural effects. Section 4 situates the work in context and clarifies its limitations.

## 2 Methods

### 2.2 Dataset

The primary training and evaluation cohort is the GDSC1 panel of Iorio and colleagues [9]. After intersecting on cell lines for which we have paired DNA methylation, gene expression, and drug response measurements, the dataset contains 987 cancer cell lines spanning 38 GDSC histology categories and 13 anatomical primary sites. Drug response labels are ln(IC_50_) values from GDSC1 [11, 12]. We retained 375 compounds for which at least 100 cell lines in our cohort have IC_50_ measurements, to ensure each drug has enough observations to be modelable. The resulting label matrix is sparse, with 295,641 valid (cell, drug) pairs realized out of 337,875 nominal (87.5% fill). Of these, approximately 246,000 pairs enter random-5fold evaluation (∼49,000 per test fold; see Supplementary Table S7); the remainder are dropped at training time by additional filters for per-fold feature availability.

Methylation comes from Gene Expression Omnibus (GEO) accession GSE68379 on the Illumina HumanMethylation450K platform [13, 14]. CpGPT preprocessing drops three probes as unmappable to its internal coordinate metadata, leaving 9,997. We retain the top 10,000 CpGs by variance across the cohort. The variance ranking is computed once on all cells but is unsupervised with respect to IC_50_.

Gene expression is GDSC Robust Multi-array Average (RMA)- normalized microarray data [15]. We keep the 5,000 most variable genes and transform them per training fold by standardization followed by principal components analysis (128 components). The per-fold protocol is the post-audit revision of an earlier global-PCA pipeline; we ablate the original in Section 3.

The PRISM Repurposing Secondary Screen [16] is used for external cross-screen validation only. DepMap 24Q4 binary somatic mutation calls (DepMap Portal release 24Q4, depmap.org/portal/data_page/?tab=allData; 880 cells, 1,314 genes) are used only in the negative-result sparse-bio ablation and are absent from the headline pipeline.

### 2.2 Evaluation protocol

We report three split types, each with multi-seed validation across initialization seeds 42, 7, and 123.

*Random 5-fold*: standard 80/20 random partitioning of (cell, drug) pairs. The same cell can appear in train and test with different drugs. We use this split to compare with published reports of model performances on random folds.

*Histology-blind (38 folds)*: leave-one-tissue-out. Each fold withholds all cell lines belonging to one GDSC histology category, with folds of fewer than five cell lines skipped. The model must generalize to cancer types it has never seen during training.

*Site-blind (13 folds)*: leave-one-anatomical-site-out. Each fold withholds all cells from one primary site. Site is coarser than histology, so held-out folds typically contain multiple histologies of the same tissue.

Within each train fold we hold the last 10 percent of training cells, cell-level not pair-level, as a deterministic validation carve-out. We used validation MSE as our early-stopping signal and the fit target for the *α*-ensemble coefficient. We ensure the test fold is never observed during training or model selection.

Per-seed metrics are means across the fold-level test R^2^ values. Cross-seed metrics are the mean of per-seed means. Across-seed standard deviation is consistently smaller than within-seed across-fold standard deviation by roughly an order of magnitude (Supplementary Table S7), which we read as evidence of stable architecture behavior under seed variation.

### 2.3 Foundation-model embeddings

*CpGPT.* CpGPT is a 100M-parameter transformer pretrained on approximately 150,000 methylation samples from 2,042 GEO studies [17], and we use it frozen to produce a 512-dimensional cell embedding per sample. Per private correspondence with the lead CpGPT author (April 2026), GSE68379 was not included in the CpGCorpus pretraining set, so there is no foundation-model contamination of our evaluation cohort.

*CLAMP.* CLAMP is a contrastive language-molecule pretraining model that produces 768-dimensional drug embeddings from SMILES inputs [18, 19], and we use it frozen. CLAMP covers all 429 GDSC1 compounds with parseable SMILES, of which the 375 screened against at least 100 cell lines form the prediction-task evaluation set, while the full 429-drug CLAMP-covered set is used only for FiLM *γ*-vector extraction in Section 3, where the prediction-task threshold does not apply. For compounds with multiple SMILES variants we use the canonical SMILES representation.

*Gene expression PCA.* Standardization and PCA-128 are both fit on training cells only within each fold. The PCA loadings (5,000 by 128) are also used as gene-node feature augmentations in the GNN.

### 2.4 Architecture I

Architecture I, illustrated in Figure 1, has three components.

**Figure 1:**
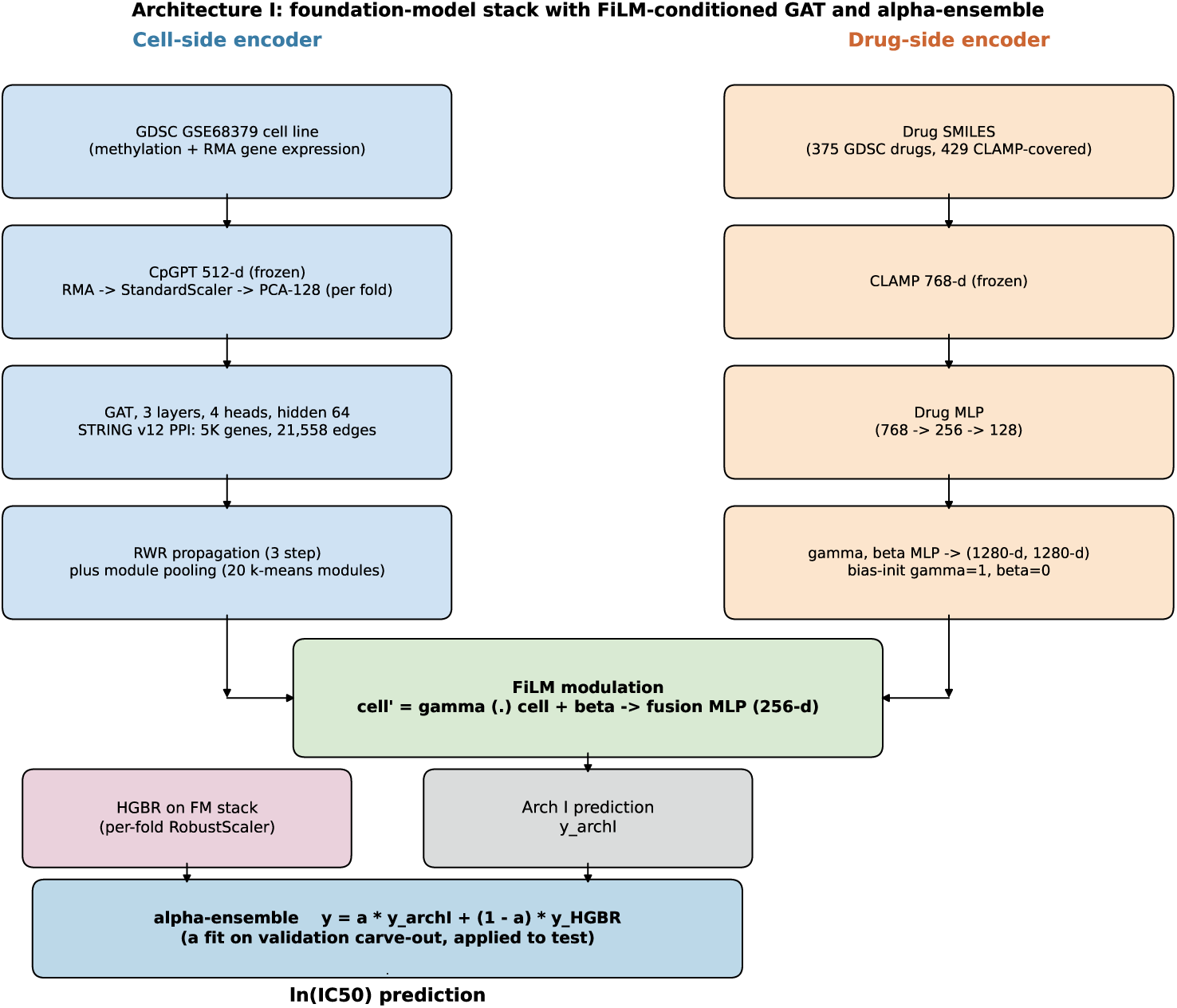
Architecture I and its components. Cell-side encoder (left, blue): frozen CpGPT plus per-fold StandardScaler and PCA-128 of gene expression, feeding a 3-layer 4-head GAT on the STRING v12 PPI graph (5,000 gene nodes, 21,558 high-confidence edges), with random-walk-with-restart propagation and 20-module pooling. Drug-side encoder (right, orange): frozen CLAMP embedding through a small MLP, with a *γ,β* head producing the FiLM modulation parameters. Fusion (center, green): **c**^′^ = *γ* ⊙**c** + *β*, then a fusion MLP to the Arch I prediction. HGBR baseline (lower left, pink) trained on the concatenated foundation-model feature vector with per-fold RobustScaler. *α*-ensemble fit on the validation carve-out, applied to test.

The cell encoder is a 3-layer, 4-head graph attention network [20] on the 5,000 gene nodes connected by 21,558 high-confidence STRING v12 edges [21]. Each node carries a 129-dimensional feature: the cell’s expression value for that gene (1-d) concatenated with the gene’s 128-d PCA loading vector. Hidden dimension is 64, and the final-layer output is pooled into 20 functional modules identified by k-means clustering on the PCA loadings. The drug encoder is a small MLP (768 to 256 to 128) over the CLAMP embedding. Fusion is by Feature-wise Linear Modulation [22]. A second small head produces drug-conditional scaling and shifting parameters *γ* and *β* (each 1,280-dimensional, matching the pooled cell representation). The modulated cell representation is **c**^′^ = *γ* ⊙**c** + *β*, where ⊙ denotes element-wise product. Both *γ* and *β* are bias-initialized to leave the FiLM block at identity (*γ* = 1, *β* = 0), so it can only help during training, never hurt at initialization. We also apply a 3-step random walk with restart [23, 24] on the PPI graph to produce a propagated cell representation concatenated with the FiLM-modulated one before the final fusion MLP.

During training we apply a similarity-augmentation regularizer that, for each batch, replaces a fraction of the cell-side input with a convex combination of its five nearest neighbors in expression space (mixing weight 0.5 on the original cell, *α* = 0.5). The neighbor graph is constructed per training fold over training cells only. This is a regularizer only and does not change inference-time predictions.

Training uses Adam at learning rate 10^−3^, plateau LR scheduling (factor 0.5, patience 10), batch size 2,048, and patience-30 early stopping on validation MSE. We note that mixed-precision bf16 autocast on A100 GPUs introduces approximately 10^−4^ R^2^ nondeterminism between runs of the same configuration.

### 2.5 HGBR baseline and *α*-ensemble

The HGBR baseline (scikit-learn’s `HistGradientBoostingRegressor` [25, 26]) uses the concatenated foundation-model feature vector of 1,408 dimensions, with per-fold RobustScaler on the cell-side features and fixed hyperparameters across all folds and seeds (`max_iter=200, max_depth=6, learning_rate`=0.1), and we apply no test-set hyperparameter selection.

The *α*-ensemble is fit per fold on the validation carve-out only. We minimize validation MSE for 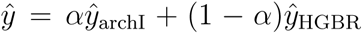 by closed-form onedimensional least squares, then apply the fitted *α* to the test fold. Across hard splits the fitted *α* falls between 0.58 and 0.61, comfortably inside the interval [0.2,0.8] we pre-specified as the gate for evidence that both component models contribute non-trivially to the ensemble.

### 2.6 Leakage discipline

We verify the pipeline against the six leakage modes catalogued by Asiaee [1]. Per-mode evidence is in Supplementary Table S1.

*M1 (supervised feature screening on full data).* The CpG variance filter is unsupervised with respect to IC_50_ so there is no ‘contamination’ in this mode.

*M2 (unsupervised preprocessing on full data).* StandardScaler, PCA-128, and the HGBR RobustScaler are all fit per training fold. An earlier version of the pipeline fit gene-expression PCA on the union of train and test cells; we retain that as the comparator for the per-fold ablation in Section 3 and show its impact is below noise on hard splits.

*M3 (test-set hyperparameter or model selection).* Hyperparameters are fixed across folds, and early stopping uses validation MSE only.

*M4 (auxiliary encoder pretraining contamination).* CpGPT’s pretraining corpus was confirmed disjoint from GSE68379 by author correspondence. CLAMP was pretrained on independent ligand-language corpora unrelated to drug response.

*M5 (improper cross-validation aggregation).* We report mean-of-per-fold-R^2^, not pooled-prediction-then-scored R^2^.

*M6 (post-hoc test-metric selection).* No best-test-epoch selection. Reported metrics are from the final checkpoint selected on validation MSE.

### 2.7 DummyDrugAvg baseline

For each (split, seed, fold) the DummyDrugAvg predictor computes the mean ln(IC_50_) per drug across the training cells of that fold, and predicts that drug-mean for every test (cell, drug) pair regardless of cell identity. Drugs unseen in train fall back to the global training mean. We report this baseline as a row in Table 1 on the recommendation of [2] and [3], who showed that the cell-aware lift above the drug-mean floor is the meaningful comparison on cell-line drug response benchmarks.

**Table 1:**
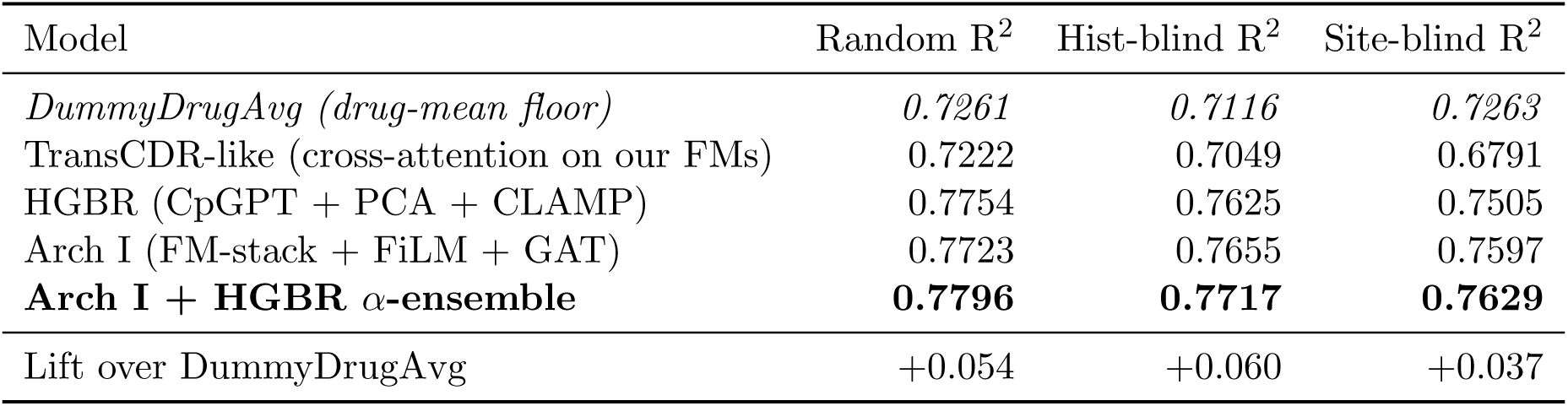
Headline performance across three evaluation splits. R^2^ values are means across folds, multi-seed across initialization seeds 42, 7, and 123. The cell-aware lift over the DummyDrugAvg baseline (italic row) is the load-bearing comparison. See Methods for split definitions.

## 3 Results

### 3.1 Headline performance and the drug-mean floor

Table 1 reports test R^2^ and pair-pooled Pearson correlation across five model classes and three split types. Figure 2 visualizes the same numbers. The Arch I + HGBR *α*-ensemble reaches R^2^ of 0.7796 on random 5-fold, 0.7717 on histology-blind, and 0.7629 on site-blind. Pair-pooled Pearson correlations are approximately 0.882, 0.879, and 0.873 across the three splits, placing the model in the 0.85 to 0.90 range that recent leakage-aware re-evaluations of published methods report [4, 3, 5].

**Figure 2:**
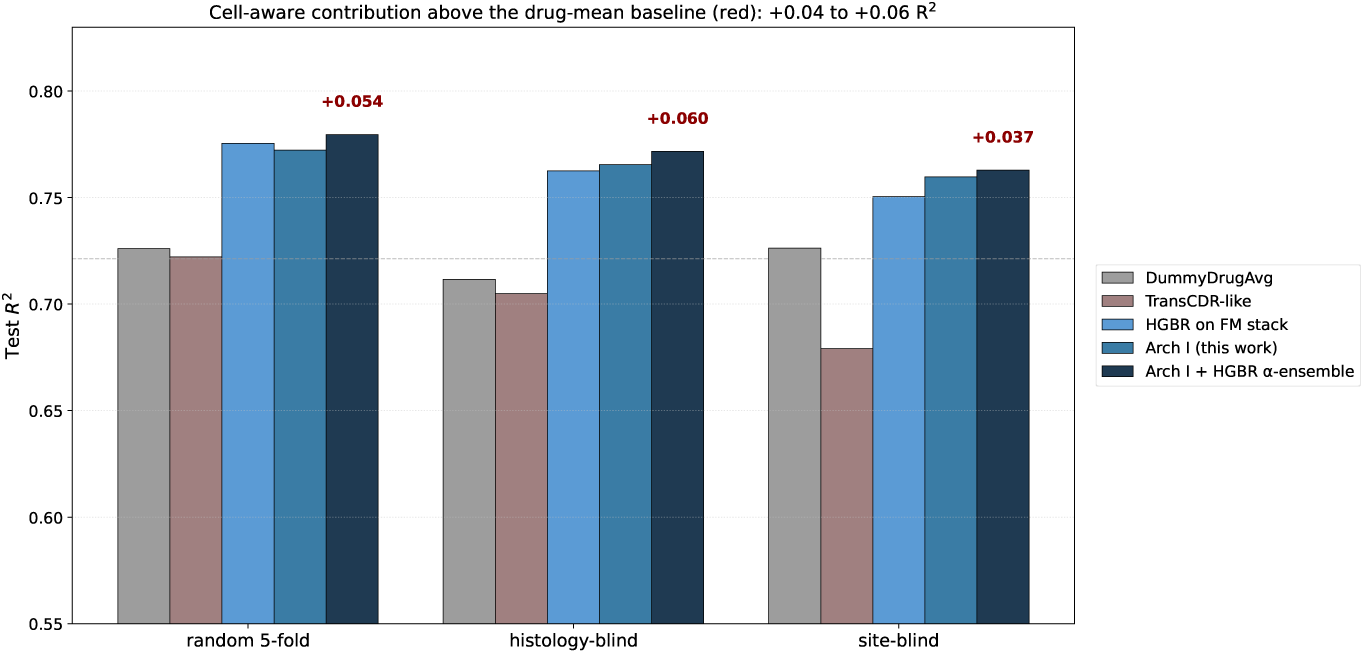
Headline performance across three evaluation splits. Test R^2^ for five model classes on random 5-fold, histology-blind, and site-blind. The DummyDrugAvg baseline (gray) sets the drug-identity floor at R^2^ around 0.71 to 0.73. Red annotations show the lift of the Arch I + HGBR *α*-ensemble over DummyDrugAvg: +0.054 random, +0.060 histology-blind, +0.037 site-blind. Multi-seed across seeds 42, 7, and 123.

The DummyDrugAvg baseline reaches R^2^ of 0.7261, 0.7116, and 0.7263, with pair-pooled Pearson correlations of 0.852, 0.850, and 0.844 across the three splits. This replicates the finding of [2]: a predictor that knows only drug identity, with no cell-side input at all, sits within 0.09 Pearson of the published 0.94 state-of-the-art numbers. The cell-aware lift above this floor is 0.054, 0.060, and 0.037 R^2^ across the three splits. We argue that this lift, not the absolute R^2^ or Pearson, is the more informative comparison for a cell-line drug response method to be evaluated against.

### 3.2 The representation jump explains most of the gain

Figure 3 decomposes the headline performance across five phases of the project. Flat-feature baselines (random forests, gradient boosting, Lasso, and generalized additive models, all trained on the raw 10,000 CpG plus 5,000 gene matrix) reach R^2^ of 0.05 to 0.10 on the random split but collapse to negative R^2^ on both hard splits. The histology-blind R^2^ from those baselines is around −0.06, meaning the flat-feature models predict worse than the global IC_50_ mean once tissue is held out.

**Figure 3:**
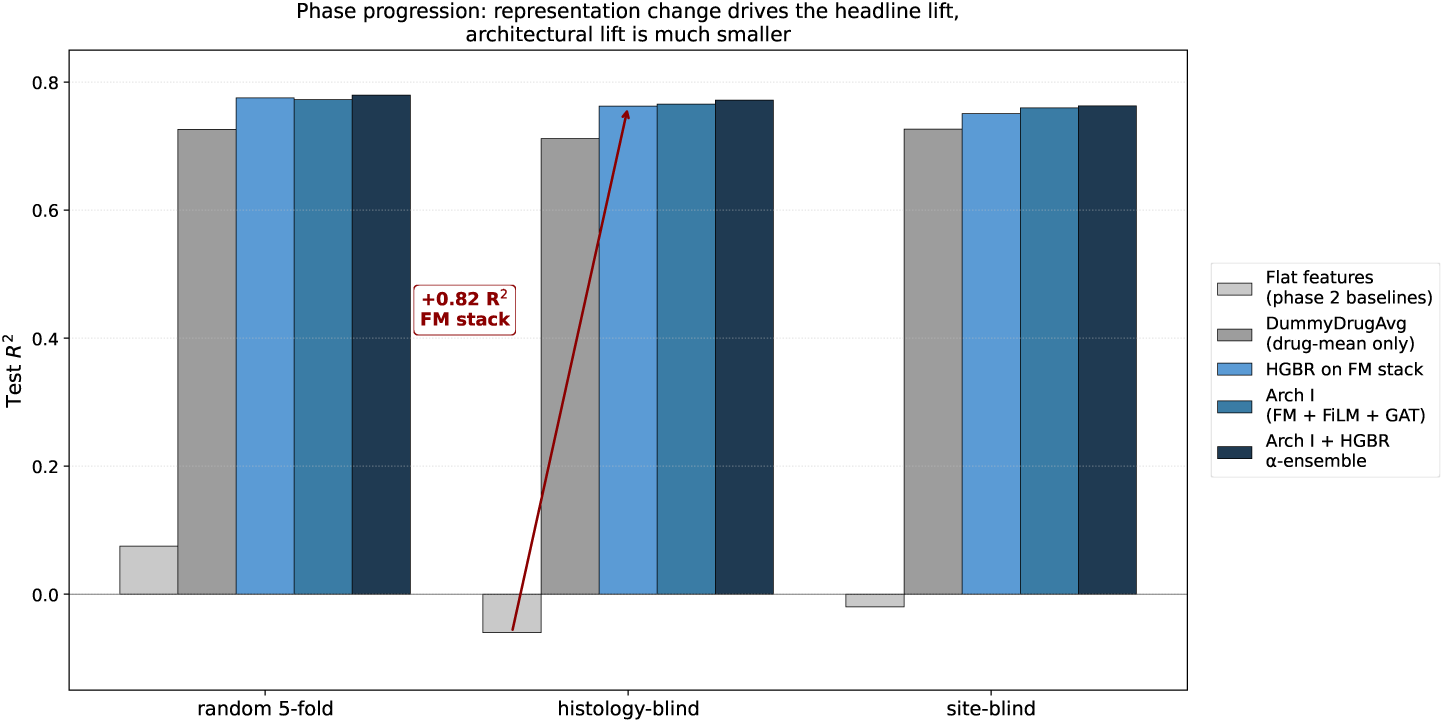
Phase progression on three evaluation splits. Test R^2^ for flat features (RF / gradient boosting / Lasso / GAM on raw 10,000 CpGs + 5,000 genes), DummyDrugAvg, HGBR on foundation-model stack, Arch I, and Arch I + HGBR *α*-ensemble. The histology-blind jump from −0.06 to +0.7625 is achieved by representation change alone. The architectural lift on top is roughly an order of magnitude smaller.

Replacing the raw input with foundation-model embeddings, while keeping the model class fixed at histogram-based gradient boosting, lifts histology-blind R^2^ to 0.7625. The single-step gain is roughly +0.82 R^2^ on histology-blind, achieved by representation change alone, before any architectural development. The architectural lift on top of HGBR-on-FM-stack, that is, the addition of Arch I and the *α*-ensemble, adds a further +0.009 R^2^ on histology-blind. The architectural contribution is significant, but its magnitude is roughly an order of magnitude smaller than the representation contribution. The paper does not headline architecture as a result.

### 3.3 Cross-architecture residual correlation puts a number on the tissue-lineage ceiling

That the predictive signal in cell-line drug response is dominated by tissue lineage is not a finding of this paper. It is established in the original GDSC landscape analysis [9], re-articulated in cross-method assessments [4], and confirmed in recent generative-framework work [10]. What we contribute is a number for the ceiling.

For each (split, seed, fold) we compute the per-pair residual *r_M_* = *ŷ_M_* − *y*_true_ for both Arch I and the HGBR baseline on the test fold. The Pearson correlation between Arch I and HGBR residuals on histology-blind hard splits is *ρ* = +0.939, averaged across all 38 folds and 3 seeds. The same statistic on site-blind is +0.949, and on random 5-fold is +0.950. All three are well above 0.9. Two radically different architectures on the same feature stack agree on most of their residual error.

Figure 4 shows the residual scatter on the largest histology-blind fold (lung NSCLC adenocarcinoma, *n* = 16,784 aligned test pairs). The cross-architecture correlation on this fold is *ρ* = +0.948. The inset gives the comparator that makes this number meaningful: the Pearson correlation between residuals from two independently initialized Arch I models on the same fold is *ρ* = +0.985, or +0.9736 ± 0.0109 aggregated across all 114 fold-pairs on histology-blind. The 0.97 versus 0.94 gap on histology-blind, roughly 0.03, is the architecture-specific signal. The rest is shared lineage variance encoded in the foundation-model features.

**Figure 4:**
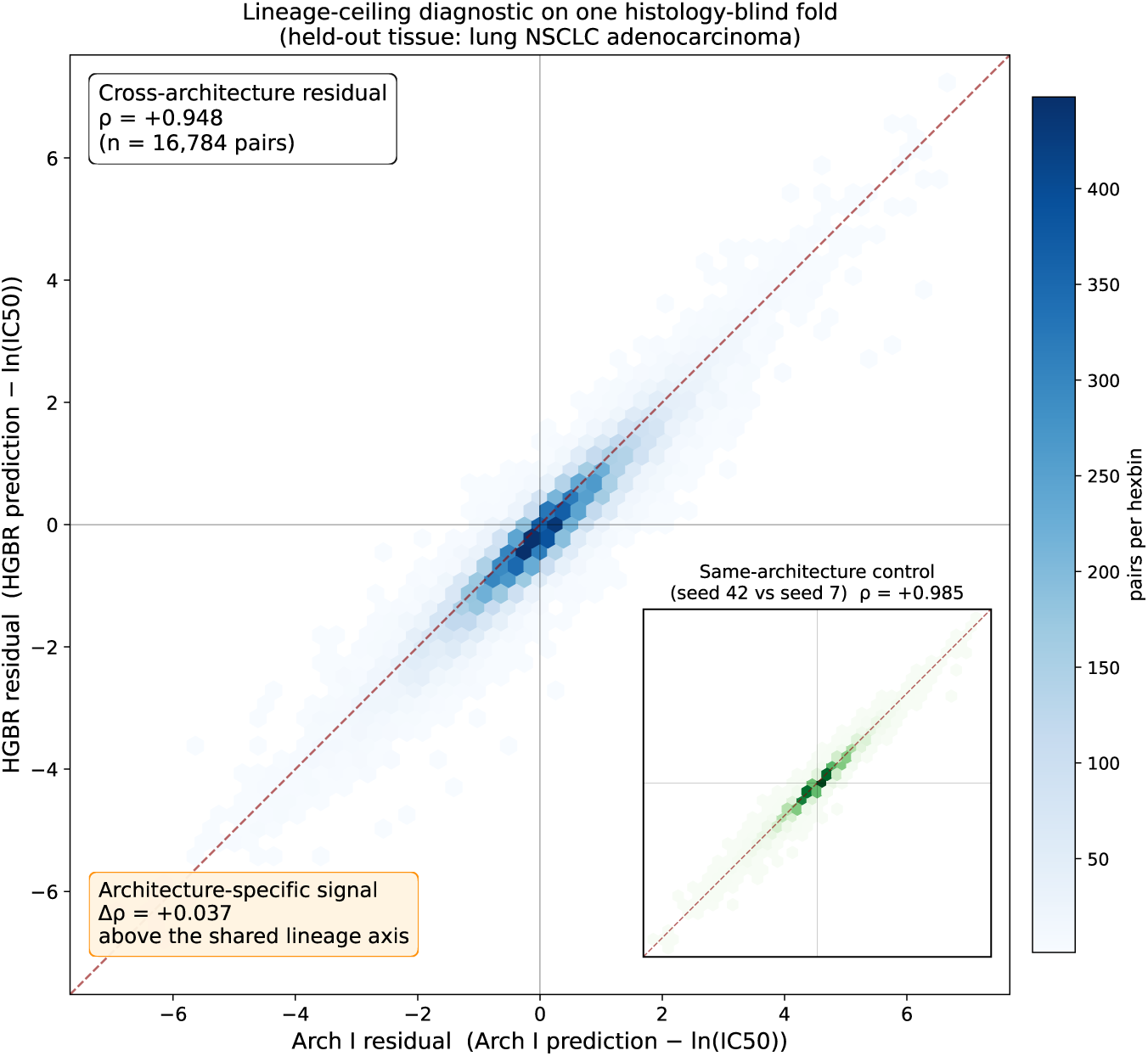
Cross-architecture residual correlation quantifies the tissue-lineage ceiling. Test-set residuals from Arch I (seed 42) and the HGBR baseline on a representative histology-blind fold (held-out tissue: lung NSCLC adenocarcinoma, *n* = 16,784). Cross-architecture Pearson correlation *ρ* = +0.948 on this fold, with aggregate *ρ* = +0.939 across all 38 hist-blind folds and 3 seeds. Inset: same-architecture control, residuals from two independently initialized Arch I models (seed 42 vs seed 7) on the identical fold, *ρ* = +0.985 (aggregate +0.974). The 0.037 gap on this fold (0.034 in aggregate) is the architecture-specific signal above the shared tissue-lineage axis.

We propose this metric, cross-architecture residual *ρ* calibrated against a same-architecture-different-initialization control, as a quantitative diagnostic for the saturation of a feature stack with respect to a particular pair of model classes. It is cheap to compute: no retraining is required if per-fold predictions are stored for two model classes and two initializations of one of them. The intuition is that values close to the same-architecture ceiling indicate the architectures recover similar predictive structure from the shared inputs, while a wider gap indicates the architectures are extracting decorrelated signal from the same features.

### 3.4 Curated biological priors do not improve prediction

We tested whether curated mutation, methylation, and drug-target-expression channels provide signal beyond the foundation-model stack. The augmented Arch I has an additional SparseBioHead, a 64 to 32 MLP whose last layer is zero-initialized so the augmented model is identical to baseline Arch I at initialization. The head’s 32-d output is concatenated into the fusion MLP. Three channels were tested:

- *Driver mutations.* 17 GDSC-relevant driver genes from DepMap 24Q4 binary calls, restricted to those at population frequency at least 5 percent (these are a different 17-gene panel from the canonical-driver set used in the CpGPT linear-probing analysis in Supplementary Section S10; overlap is partial).
- *Pharmacogenomic CpG panel.* 13 canonical methylation probes covering MGMT, MLH1, BRCA1, CDKN2A, GSTP1, and RASSF1A.
- *Drug-target expression.* 69 genes encoding direct targets of GDSC drugs, z-scored expression.

The mutations channel alone produces ΔR^2^ = −0.0036 on random 5-fold compared to baseline Arch I (across-seed std 0.0019). The all-three-channels-combined configuration produces ΔR^2^ = −0.0146 (across-seed std 0.0007). The deficits are small but reproducibly negative. The combined-channel deficit is larger than the mutations-only deficit, consistent with the interpretation that adding more sparse-bio columns through a shared head uses gradient capacity without bringing decorrelated signal.

We interpret the result as feature-stack saturation rather than as a property of the SparseBioHead design. CpGPT embeddings linearly encode tissue at 9.1 times above chance and histology at 21.1 times above chance, and curated mutation, methylation, and drug-target-expression channels are themselves substantially lineage-correlated [27, 28]. The 99 added columns mostly carry information the foundation-model embeddings already encode densely. We are not claiming that mutations are biologically irrelevant to drug response; the original GDSC analysis documented many specific mutation-drug interactions [9]. We claim only that, conditional on a CpGPT plus RMA-PCA cell representation, the curated channels in our configuration add no aggregate R^2^.

### 3.5 The global-PCA leak is non-load-bearing on hard splits

The leakage-audit flagged a residual M2 pathway in the original pipeline, comprising six `fit()` calls: a global-fit gene-expression PCA on all 987 cells, plus the similarity-augmentation *k*-nearest-neighbors graph also fit on all 987 cells (full per-mode breakdown in Supplementary Table S1). To quantify whether these vectors inflated the headline numbers, we refit the StandardScaler, PCA-128, and the sim-aug *k*-nearest-neighbors graph on training cells only for one representative fold per split. The three folds were random 5fold fold 0, histology-blind large intestine, and site-blind lung. Arch I was retrained against the per-fold-clean preprocessing.

The per-fold-clean test R^2^ versus the global-PCA baseline:

- Random 5-fold, fold 0: 0.7515 (per-fold-clean) vs 0.7491 (global, sparse-bio-mutations seed 42 run used as the closest matched-seed global-PCA baseline since the vanilla seed-42 random-5fold run did not store per-fold predictions), Δ = +0.0024.
- Histology-blind, large intestine: 0.7027 vs 0.7039 (global, seed 42), Δ = −0.0012.
- Site-blind, lung: 0.7595 vs 0.7563 (global, seed 42), Δ = +0.0032.

All three deltas are within ±0.005 R^2^, well below any meaningful effect size. Two are positive (the per-fold-clean variant is slightly better), and one is negative. The M2 leakage pathway is empirically non-load-bearing on hard splits, so the headline numbers in Table 1 stand.

### 3.6 FiLM *γ* structure is cross-seed reproducible at the drug-grouping level, not at the pathway level

We extracted the FiLM *γ* and *β* vectors for all 429 drugs across three independently trained Arch I checkpoints (seeds 42, 7, 123) and computed pathway-pair cosine similarity matrices per seed, grouping drugs by GDSC TARGET PATHWAY annotation (23 pathways with at least 3 drug members, yielding 253 pathway pairs). Cross-seed Pearson correlations on the upper-triangular entries are 0.528 (seeds 42 vs 7), 0.675 (seeds 42 vs 123), and 0.560 (seeds 7 vs 123).

To test whether this cross-seed agreement reflects pathway-specific structure or any consistent drug grouping, we ran a 1,000-permutation null in which drug-to-pathway assignments were shuffled before recomputing the pathway-pair similarity matrices for each seed. Under the shuffled labeling the null cross-seed correlations have mean values of 0.541, 0.601, and 0.517 (standard deviations roughly 0.10 for each pair). The observed correlations fall within the null distribution, with empirical one-sided p-values of 0.565, 0.230, and 0.367. The cross-seed reproducibility we see is therefore a property of any consistent drug grouping on these embeddings.

*γ*-space structure in the FiLM analysis is reproducible across random seeds. It is misleading to claim the structure reflects biological pathway grouping when an equally consistent random grouping yields the same magnitude of cross-seed agreement. We retain the per-seed *γ* heatmaps and the mean-centered pathway-pair matrix as supplementary figures with this hedge in their captions, rather than as a headline biological interpretation result.

### 3.7 Multi-seed stability

Across-seed standard deviations on R^2^ are approximately an order of magnitude smaller than within-seed across-fold standard deviations: on random 5-fold the seed-std is 0.0006 with fold-std around 0.011, on histology-blind the seed-std is also 0.0006, and on site-blind the seed-std rises to 0.0058 because the 13 sites differ more substantially in test-set composition than the 38 histologies do. The full per-seed per-fold matrix is reported in Supplementary Table S7, confirming that the headline numbers are not artifacts of the seed-42 draw.

### 3.8 Cross-screen validation against PRISM matches the measurement reproducibility ceiling

We froze one trained Arch I checkpoint (seed 42, random 5-fold fold 0) and predicted ln(IC_50_) for 39,682 (cell, drug) pairs from the PRISM Repurposing Secondary Screen [16], without any retraining or fine-tuning on PRISM data. Of these pairs, 26,714 are also in GDSC and provide the cross-screen measurement-reproducibility ceiling.

The model’s global Pearson correlation against PRISM is 0.225 and its Spearman correlation is 0.627. The GDSC-to-PRISM measurement ceiling, computed by treating the GDSC IC_50_ values themselves as predictions of the PRISM values on the shared (cell, drug) pairs, is 0.207 Pearson and 0.635 Spearman. The model matches Spearman within 0.01 and slightly exceeds the Pearson ceiling. Figure 5 reports both metrics alongside the per-drug Pearson distribution.

**Figure 5:**
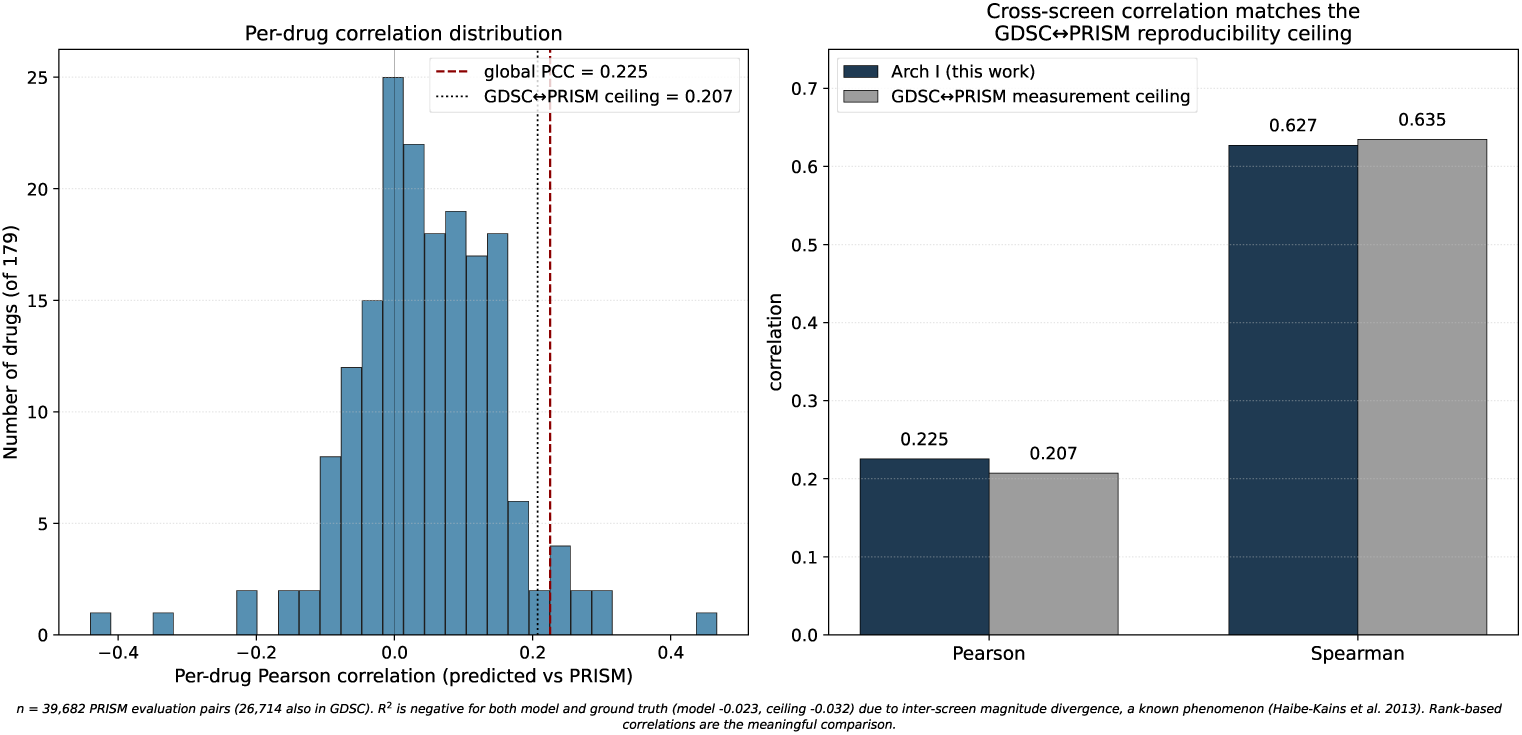
Cross-screen validation against PRISM. Left: distribution of per-drug Pearson correlations across 179 shared drugs (*n*_drug_ ≥ 5). Right: model versus GDSC-to-PRISM measurement reproducibility ceiling on Pearson and Spearman correlation. Model Spearman 0.627 is within 0.01 of ceiling 0.635; model Pearson 0.225 slightly exceeds ceiling 0.207. R^2^ is negative for both the model and the GDSC ground truth, reflecting inter-screen magnitude divergence [29]; rank-based correlations are the appropriate cross-screen comparison.

R^2^ is negative for both the model (−0.023) and the GDSC ground truth (−0.032), as expected from inter-screen magnitude divergence: GDSC and PRISM use different dose ranges and viability readouts, so the absolute IC_50_ magnitudes do not align even when both measurements are correct [29]. Rank-based correlations are the appropriate cross-screen comparison, and on those metrics the model captures the transferable rank signal between the two screens up to the measurement reproducibility ceiling.

### 3.9 TransCDR-like architectural control

To isolate the contribution of architecture choice from input choice, we reimplemented the TransCDR fusion mechanism [30] on identical foundation-model inputs and trained it under the same multi-seed and three-split protocol. The TransCDR-like comparator reaches R^2^ of 0.7222, 0.7049, and 0.6791 across random, histology-blind, and site-blind. Arch I reaches 0.7723, 0.7655, and 0.7597 under the same protocol. The margin is 5 to 8 R^2^ points and widens on harder splits.

This comparison is architecture-only on clean inputs. The published TransCDR pipeline uses different inputs (z-scored RNA and methylation, mutation, and GIN drug graphs) and reports Pearson 0.94 on warm-start random splits. We are not claiming to beat the published TransCDR under leakage-aware re-evaluation. The public TransCDR training code reads z-scored matrices computed on the full cell-line set before applying crossvalidation, which is Asiaee mode M2 leakage, and would require a substantive rewrite to test under matched discipline.

## 4 Discussion

Tissue lineage dominates the predictive signal in cell-line drug response. That has been established for a decade [9], replicated in cross-method assessments [4], and re-articulated under generative frameworks [10]. The Connectivity Map / L1000 work tells the same story for transcriptional perturbation response more broadly [31].

On histology-blind splits, where we have the most fold-pairs (114 across 38 folds and 3 seed pairs), the cross-architecture residual correlation is *ρ* = 0.939 against a same-architecture-different-initialization ceiling of 0.9736±0.0109, a Δ = +0.0346 gap that clears our pre-specified threshold of +0.03 for nontrivial architecture-specific signal. On site-blind and random 5-fold the gaps are smaller (+0.024 and +0.020) and below threshold (Supplementary Table S4); the histology-blind PASS is the load-bearing measurement. Either way, the headroom that depends on architecture choice given the current feature stack is small. We expect that heavier architectures, including graph transformers, mixture-of-experts fusion blocks, and larger language-model fusion heads, would not move this number substantially given the present cell representation, although we have not directly tested them under our protocol. The implication is that the bottleneck appears to lie in the representation rather than in the architecture. Promising directions for moving past this ceiling include cell features that better decorrelate from the lineage axis, such as spatial or single-cell-resolved methylation atlases, multi-tissue perturbation profiles, or different prediction targets such as per-cell residuals after a tissue-mean baseline.

### 4.1 Why curated biological priors did not help

The mutation, methylation, and drug-target-expression channels carry significant biological information. Driver mutations have well-documented and often clinically actionable drug-response interactions [9, 32, 33]. Why, then, do they fail to improve aggregate R^2^ once foundation-model embeddings are in place?

The cleanest answer is feature-stack saturation along the lineage axis. CpGPT linearly encodes tissue at 9.1 times above chance and histology at 21.1 times above chance. Mutation frequencies are strongly tissue-stratified [27]. Pharmacogenomic methylation loci segregate by histology [28, 32]. Drug-target gene expression varies systematically across tissues. The 99 sparse-bio columns therefore project largely onto the same low-rank subspace the foundation-model embeddings already encode. The 32-dimensional SparseBioHead output consumes gradient capacity without bringing decorrelated signal.

This is a statement about aggregate R^2^ on the GDSC panel. It is not a statement that mutations are biologically irrelevant. A more targeted use, per-drug stratified prediction, biomarker discovery for a specific mutation-drug interaction, or fine-tuning to a panel of compounds with known mutation-driven sensitivity, might still benefit from these channels even under our feature stack. The negative result here is conditional on a specific evaluation regime, and we make no claim about other regimes.

### 4.2 What the leakage-clean window means for the field

Of the 32 audited methods in the public companion roster maintained by Asiaee and colleagues at the time of writing [1], six are certified clean, and none of them combines a methylation foundation model with a drug foundation model under multi-omics input and leave-tissue-out hard splits with multi-seed validation; the intersection is, as of May 2026, an empty cell. We expect that leakage-aware evaluation will increasingly become a standard expectation in this area as the audit literature is taken up by reviewers.

Pillbox fills that cell, though we do not interpret this as conferring special standing on our work, and we expect the cell to fill in over the next year as the leakage-aware reframing spreads. We do propose the present pipeline as a template: a per-fold preprocessing discipline that closes M2 by construction, an external corpus disjoint check by author correspondence for M4, a cross-architecture residual diagnostic to test saturation of the chosen feature stack, and an honest hedge on interpretability claims that do not survive a permutation null.

### 4.3 Limitations

There is a non-trivial clinical translation gap: we evaluate end-to-end on cell-line data with cross-screen validation against PRISM, but we do not have patient-derived organoid, patient-derived xenograft, or clinical-trial data, and recent translational work in this area has reached a higher bar than ours [34, 35]. A natural extension of the present pipeline would be to use the Arch I + HGBR ensemble as a pretrained stage and fine-tune on PDO or PDX data before clinical inference, but we have not done this.

The graph attention network used in our cell encoder is a 2018 architecture, which has proven empirically to be somewhat outdated. Graph transformers and long-range message-passing variants are now the medium-tier standard. A graph-transformer ablation under our protocol is realistic and expected future work. Given the lineage-ceiling diagnostic we report, we do not expect such an ablation to move hard-split R^2^ by more than roughly the 0.03 architectural headroom.

The FiLM *γ*-space pathway-pair interpretation does not survive a permutation null. The structure in *γ*-space exists and is cross-seed reproducible, but it is not pathway-driven. Earlier internal analyses identified specific cross-pathway pairs (DNA Replication with Mitosis, EGFR with WNT, and so on) with biologically plausible interpretations. We do not lead with these results because we cannot show they exceed what an arbitrary drug grouping would produce on the same *γ* vectors. The honest framing is that FiLM *γ*-space is reproducible across seeds, the specific pathway interpretation is not distinguishable from arbitrary groupings, and we leave the biology to further work with a pathway-survivable test.

Mixed-precision bf16 autocast introduces approximately 10^−4^ R^2^ nondeterminism, and we did not enable deterministic CUDA flags; reported numbers are reproducible within this tolerance, and absolute differences smaller than 10^−3^ R^2^ between configurations should be treated as within noise.

Finally, all headline results are on GDSC1. PRISM cross-screen validation provides external evidence against a different screen on largely overlapping cell lines, but we have not retrained or fine-tuned on PRISM. Datasets used by the broader literature (CCLE, gCSI, CTRPv2) are absent from our evaluation. The IMPROVE benchmark [5] provides matched preprocessing across all five and would be the natural cross-dataset extension. We expect the cross-dataset generalization picture for our method to look similar to other clean methods evaluated under IMPROVE: substantial degradation cross-dataset, consistent with what [4] and [3] report.

## 5 Conclusion

Cancer drug response prediction on cell-line panels is in a period of methodological reassessment. The published accuracy numbers in this area appear to be substantially shaped by preprocessing protocol and by drug-identity variance, rather than by cell-aware predictive signal alone. Once leakage is audited away and the floor is set at drug-mean prediction, the cell-aware contribution we observe is small, in the 0.04 to 0.06 R^2^ range, and similar in magnitude to what other recent leakage-aware re-evaluations report. Our specific contribution is a leakage-aware predictor combining foundation-model embeddings under multi-seed hard splits, paired with a residual-correlation diagnostic that puts a number on the tissue-lineage ceiling for our feature stack.

## Ethics Statement

No ethical approval was required for this work. All data analyzed are publicly available molecular and pharmacological measurements of established cancer cell lines (GDSC1, PRISM, DepMap, GEO accession GSE68379); no human or animal subjects, patient data, or identifiable specimens were involved.

## Acknowledgements

This work used the Advanced Cyberinfrastructure Coordination Ecosystem: Services & Support (ACCESS), supported by the National Science Foundation (NSF).

This material is also based upon work supported by the NSF Graduate Research Fellowship Program under Grant No. DGE-2139757. Any opinions, findings, and conclusions or recommendations expressed in this material are those of the authors and do not necessarily reflect the views of the NSF.

## Author Contributions

J.H. designed the study, led the analysis, and wrote the manuscript. E.J. contributed to the GNN architecture experiments and the FiLM interpretability analysis. S.S., A.G., and D.A. contributed to data curation, baseline bench-marking, and the leakage audit. J.H.J. contributed to the foundation-model embedding pipeline and cross-screen validation. H.J.R. secured computational resources, and contributed to the biological-priors ablation and the clinical-translation framing. All authors reviewed and approved the final manuscript.

## Supplementary Information

### S1. Asiaee M1–M6 leakage audit (Supplementary Table S1)

The audit covers every `fit()` and `fit_transform()` call in the pill-box codebase outside of visualization and archived scripts. Each call is classified as M1–M6 clean, leaky against a specific Asiaee mode, or non-training (N/A). The HGBR pipeline (`run_xgb_hard_splits.py`, `run_xgb_mutation_ablation.py`; filename retains the historical `xgb` prefix for the legacy gradient-boosting baseline, but the imported class is scikit-learn’s `HistGradientBoostingRegressor`) is per-fold clean throughout: `RobustScaler` is fit on *X*_tr_ only and `transform` is applied to *X*_va_ and *X*_te_. The GNN pipeline has six pre-audit M2 vectors: the gene-expression `StandardScaler` + PCA-128 fit on the full 987-cell matrix (`scripts/encode_gene_expr.py:173-177`, `scripts/encode_gene_expr.py:363-366`), and the similarity-augmentation `NearestNeighbors.fit(expr_np)` over all 987 cells used at train time (`run_hard_splits.py:113`, `train_gnn.py:200`, `scripts/dump_arch_i_predictions.py:92`). The per-fold PCA ablation in Section 3 and Supplementary Table S2 shows the impact of these vectors is within noise on hard splits. We do not patch them in the headline numbers reported in Section 3, but we do report the per-fold-clean ablation explicitly. Audit script: `scripts/audit_fit_calls.py`; raw output: `results/leakage_audit.json`.

### S2. Per-fold PCA ablation (Supplementary Table S2)

For one representative fold per split, we refit `StandardScaler` + PCA128 + the sim-aug *k*-nearest-neighbors graph on training cells only and retrain Arch I. The fold-level test R^2^ values are 0.7515 (random 5fold/fold 0, vs 0.7491 global-PCA baseline), 0.7027 (histology blind/large intestine, vs 0.7039), and 0.7595 (site blind/lung, vs 0.7563). All three deltas are within ±0.005 R^2^; two are positive (per-fold-clean better) and one is slightly negative. The M2 leakage pathway is non-load-bearing for our headline numbers.

Build script: `scripts/per_fold_pca_ablation.py`. The `PILLBOX_TRAIN_DIR` and `PILLBOX_EMB_DIR` environment variables override the static paths in src/data/graph dataset.py so that per-fold-fitted PCA components and gene loadings replace the global ones, while all other static assets (drug lookups, split JSONs, methylation arrays) remain shared by hardlink.

**Table S1:** Asiaee M1–M6 leakage audit. Every `fit()` and `fit_transform()` call in pillbox outside visualization and archived scripts, classified as clean, leaky against a specific Asiaee mode, or non-training (N/A).

| File | Line | Mode | Disposition |
| --- | --- | --- | --- |
| probe_cpgpt_representations.py | 61 | N/A | `LabelEncoder.fit_transform` on tissue/histology labels; supervised probe target encoding |
| probe_cpgpt_representations.py | 82 | clean | Per-fold `StandardScaler` on $X_{tr}$ inside `StratifiedKFold` |
| probe_cpgpt_representations.py | 86 | clean | Per-fold supervised classifier `fit` on $X_{tr}$ |
| probe_cpgpt_representations.py | 115 | clean | Per-fold `StandardScaler` on $X_{tr}$ |
| probe_cpgpt_representations.py | 119 | clean | Per-fold supervised classifier `fit` on $X_{tr}$ |
| run_hard_splits.py | 113 | M2 | `NearestNeighbors.fit` on `expr.by_wide` (all 987 cells) for `sim_aug`; test cells in the kNN graph |
| run_transcdr_benchmark.py | 154 | clean | `RobustScaler` fit on `cell_emb[tr]` only |
| run_xgb_hard_splits.py | 113 | clean | `RobustScaler` fit on $X_{tr}$ only, then transform val/test |
| run_xgb_hard_splits.py | 123 | clean | `HistGradientBoostingRegressor.fit` on $(X_{tr}, y_{tr})$ |
| run_xgb_mutation_ablation.py | 54 | clean | `RobustScaler` fit on $X_{tr}$ only |
| run_xgb_mutation_ablation.py | 62 | clean | `HistGradientBoostingRegressor.fit` on train |
| train_gnn.py | 200 | M2 | `NearestNeighbors.fit` on `expr.by_wide` (all 987 cells) for `sim_aug`; test cells in the kNN graph |
| scripts/dump_arch_i_predictions.py | 92 | M2 | `NearestNeighbors.fit` on full `expr`; only used for inference re-loading, but matches the trained-time graph |
| scripts/encode_drugs.py | 553 | N/A | `TSNE.fit_transform`; visualization only |
| scripts/encode_drugs_clamp.py | 249 | N/A | `TSNE.fit_transform`; visualization only |
| scripts/encode_drugs_clamp.py | 277 | N/A | `PCA.fit_transform` on ECFP for 2-D viz only |
| scripts/encode_drugs_clamp.py | 285 | N/A | `PCA.fit_transform` on CLAMP for 2-D viz only |
| scripts/encode_gene_expr.py | 173 | M2 | `StandardScaler.fit_transform` on full expression matrix (all 987 cells) |
| scripts/encode_gene_expr.py | 177 | M2 | `PCA.fit_transform` on full standardized expression matrix |
| scripts/encode_gene_expr.py | 363 | M2 | `StandardScaler.fit_transform` on full expression matrix (RMT eigenvalue analysis path) |
| scripts/encode_methylation_cpgpt.py | 376 | N/A | `PCA.fit_transform` for 2-D embedding visualization only |
| src/visualization/plot_gse270494.py | 311 | N/A | `StandardScaler` for visualization only |

### S3. FiLM permutation null (Supplementary Table S3)

We extracted *γ* and *β* vectors for all 429 drugs from three independently trained Arch I checkpoints (seeds 42, 7, 123, fold 0 of random 5fold). We grouped drugs by GDSC TARGET_PATHWAY annotation (filtered to pathways with ≥ 3 drugs, yielding 23 pathway groups), centered the pathway-mean *γ* vectors within each seed, computed the 23 × 23 cosine-similarity matrix per seed, and Pearson-correlated the upper-triangular entries (*n* = 253 pathway pairs) across seed pairs.

**Table S2:** Per-fold PCA ablation. For one representative fold per split, we refit StandardScaler + PCA-128 + the sim-aug *k*-NN graph on training cells only and retrain Arch I. The global-PCA baseline is the original pipeline; for random 5fold/fold 0 the matched-seed comparator is the sparsebio-mutations seed-42 run (the vanilla seed-42 random-5fold run did not store per-fold predictions, and sparse-bio-mut Δ vs vanilla is −0.004 across seeds).

| Split | Fold | Per-fold-clean $R^2$ | Global-PCA $R^2$ | $\Delta R^2$ |
| --- | --- | --- | --- | --- |
| random_5fold | fold_0 | 0.7515 | 0.7491 | +0.0024 |
| histology_blind | large_intestine | 0.7027 | 0.7039 | $-0.0012$ |
| site_blind | lung | 0.7595 | 0.7563 | +0.0032 |

To test pathway-specificity, we shuffled the drug-to-pathway assignment uniformly at random and recomputed all three cross-seed correlations under the shuffled labeling, repeated for 1,000 permutations. Observed correlations are within the null distribution, indicating that the cross-seed reproducibility of pathway-pair similarity is a property of any consistent drug grouping rather than a property of the GDSC pathway annotation specifically.

We retain the per-seed *γ*-space heatmaps and the mean-centered pathway-pair similarity matrix as supplementary figures with hedged captions. The robust positive and negative pathway pairs reported in earlier internal analyses (DNA replication ↔ Mitosis, EGFR ↔ WNT, Cytoskeleton ↔ RTK signaling, and the corresponding anti-correlations) are not claimed as supported by cross-seed reproducibility alone; they may still reflect underlying biology but require an independent permutation-survivable test. Script: `scripts/film_pathway_permutation_null.py`.

**Table S3:** FiLM *γ*-space cross-seed agreement under a drug-to-pathway permutation null. Pearson *r* on upper-triangular pathway-pair cosine-similarity entries (*n* = 253 pairs across 23 pathways). All three observed values fall within the shuffled null distribution.

| Seed pair | Obs. $r$ | Null mean | Null std | Null p95 | $p$ (1-sided) | Verdict |
| --- | --- | --- | --- | --- | --- | --- |
| 42 vs 7 | +0.5281 | +0.5410 | 0.0993 | +0.6911 | 0.5654 | n.s. |
| 42 vs 123 | +0.6754 | +0.6009 | 0.0977 | +0.7496 | 0.2298 | n.s. |
| 7 vs 123 | +0.5598 | +0.5168 | 0.1105 | +0.6904 | 0.3666 | n.s. |

### S4. Cross-architecture residual-*ρ* null (Supplementary Table S4)

The headline biology claim in Section 3 is that cross-architecture residual correlation between Arch I and HGBR on hard splits is meaningfully below the same-architecture-different-initialization ceiling. To compute the null, we used the per-fold prediction arrays from three independently-initialized Arch I models (seeds 42, 7, 123) on the same fold definitions and computed Pearson residual correlations between all (seed-A, seed-B) pairs.

The histology-blind split yields a same-arch overall correlation of 0.9736± 0.0109 (*n* = 114 fold-pairs across 38 folds × 3 seed pairs), compared to the cross-architecture observed value of 0.939. The Δ = +0.0346 exceeds our pre-specified threshold of +0.03 for “same-arch *ρ* exceeds cross-arch *ρ* by a margin compatible with non-trivial architecture-specific signal.” The site-blind and random 5fold splits yield Δ values of +0.0241 and +0.0200 respectively, positive in direction but below the +0.03 threshold. We classify these as WEAK in the table but note that all three directions are consistent.

random 5fold has only 5 fold-pairs because only the (seed7, seed123) pair has the per-fold prediction arrays needed for the computation; the seed42 random 5fold preds were not dumped during the original multi-seed run. Script: `scripts/residual_rho_null.py`.

### S5. Architecture history (Supplementary Table S5)

We document the full progression of nine GNN architectures (A through I) plus the final *α*-ensemble, including negative results that did not enter the headline. Architectures A and B (multi-head GAT and HGT-with-CpG-nodes) had the cell encoder detached from the gradient path due to GPU memory pressure, so only the drug encoder and prediction head were actually training; both plateaued at R^2^ ≈ 0.735 random. Architectures C and D were interim drug-target-gating variants, kept in `src/models/` for reproducibility but not paper-relevant. Architecture E (module pooling + drug targeting) was the first GNN competitive with HGBR, with its site-blind PCC of 0.874 surpassing the published TransCDR cold-cell PCC of 0.864 on a generalization metric. The arch-E v2 ablation identified three positive components: similarity augmentation, inferred drug targets, and multitask auxiliary tissue prediction. Architecture F (more capacity) overfit with 880 cells and was kept as evidence for the limited-data regime. Architecture G (edge-masking + RWR propagation) showed that edge masking hurts, while RWR propagation alone provides +0.004 R^2^, indicating that mutation topology is not the bottleneck. Architecture H (cross-attention) collapsed to R^2^ ≈ 0.55 because aggressive aggregation kills the signal. Architecture I (production: GAT + FiLM + module pooling + RWR) gave R^2^ = 0.7723*/*0.7655*/*0.7597 across random / hist-blind / site-blind; with *α*-ensemble: 0.7796*/*0.7717*/*0.7629.

**Table S4:** Cross-architecture residual-*ρ* null. Same-architecture-differentinitialization residual correlation across (seed-A, seed-B) pairs of Arch I, compared to the cross-architecture (Arch I vs HGBR) residual correlation on the same fold definitions. PASS = Δ *>* +0.03 (pre-specified threshold for non-trivial architecture-specific signal); WEAK = positive but below threshold; NO DATA = required per-fold prediction arrays not on disk.

| Split | Seed pair | $n$ folds | Same-arch $\rho$ (mean ± std) | Cross-arch $\rho$ | $\Delta$ | Verdict |
| --- | --- | --- | --- | --- | --- | --- |
| random_5fold | 42-7 | n.a. | n.a. | +0.9500 | n.a. | NO DATA |
| random_5fold | 42-123 | n.a. | n.a. | +0.9500 | n.a. | NO DATA |
| random_5fold | 7-123 | 5 | $+0.9700 \pm 0.0084$ | +0.9500 | +0.0200 | WEAK |
| random_5fold | overall | 5 | $+0.9700 \pm 0.0084$ | +0.9500 | +0.0200 | WEAK |
| histology_blind | 42-7 | 38 | $+0.9731 \pm 0.0110$ | +0.9390 | +0.0341 | PASS |
| histology_blind | 42-123 | 38 | $+0.9745 \pm 0.0095$ | +0.9390 | +0.0355 | PASS |
| histology_blind | 7-123 | 38 | $+0.9733 \pm 0.0120$ | +0.9390 | +0.0343 | PASS |
| histology_blind | overall | 114 | $+0.9736 \pm 0.0109$ | +0.9390 | +0.0346 | PASS |
| site_blind | 42-7 | 13 | $+0.9757 \pm 0.0159$ | +0.9490 | +0.0267 | WEAK |
| site_blind | 42-123 | 13 | $+0.9754 \pm 0.0103$ | +0.9490 | +0.0264 | WEAK |
| site_blind | 7-123 | 13 | $+0.9682 \pm 0.0188$ | +0.9490 | +0.0192 | WEAK |
| site_blind | overall | 39 | $+0.9731 \pm 0.0158$ | +0.9490 | +0.0241 | WEAK |

The arch-A through arch-H negative results matter because they constrain the interpretation of the FiLM mechanism in Section 3: more capacity, mutation-conditioned topology, and aggressive aggregation all degrade performance under our feature stack, while FiLM-style conditioning and RWR propagation provide the small remaining lift that is then matched and slightly exceeded by *α*-ensembling with HGBR.

### S6. Asiaee 2026 leakage-clean roster comparison (Supplementary Table S6)

We list the six methods Asiaee 2026 (v2, April 2026) certified leakage-clean (Velodrome 2022, UNO/IMPROVE 2024, DeepResponse 2023, ScreenDL 2024 [7], DrVAE 2019, and MMDRP 2024 [8]) alongside one additional column for pillbox. The intersection of all eight criterion rows is occupied only by pillbox as of May 2026. We do not interpret this as a discovery; we treat it as a methodological template, and we expect this intersection to fill in over the next twelve months as the leakage-aware reframing is taken up by other methods groups.

**Table S5:** Architecture history. Progression of GNN architectures A–I plus the *α*-ensemble, including negative results that did not enter the headline. Em dashes denote configurations for which the specific split was not run. “D-###” references are entries in the project decision log.

| Arch | Year | Description | Rand R<sup>2</sup> | Hist R<sup>2</sup> | Site R<sup>2</sup> | Notes |
| --- | --- | --- | --- | --- | --- | --- |
| phase 2 baselines | 2025-Q4 | RF/GBR/Lasso/GAM on raw 10K CpGs + 5K genes | 0.05–0.10 | −0.06 | −0.02 | flat features memorize tissue shortcuts; fail on hard splits; motivates the FM pivot |
| A: GAT (PPI gene-only) | 2026-Q1 | 4-head GAT, gene nodes only | 0.735 | n.a. | n.a. | cell encoder detached from gradient due to OOM (D-062) |
| B: HGT (CpG nodes) | 2026-Q1 | heterogeneous graph transformer; CpG-to-gene regulatory edges | 0.735 | n.a. | n.a. | same detach issue; HGT params not trained (D-064) |
| C, D | 2026-Q2 | interim drug-target gating variants | n.a. | n.a. | n.a. | not paper-relevant; kept in `src/models/` |
| E: module pool + drug targeting | 2026-Q3 | GAT + 20-module pooling + target-aware reweighting | 0.747 | 0.760 | 0.758 | site-blind PCC 0.874 first beat TransCDR cold-cell PCC 0.864 (D-067) |
| E v2 winners | 2026-Q3 | sim_aug + infer_targets + multitask | n.a. | n.a. | n.a. | D-069: all three positive; FiLM later subsumes infer_targets |
| F: more capacity | 2026-Q4 | deeper/wider GAT | n.a. | n.a. | n.a. | overfits with 880 cells; limited-data regime evidence (D-074) |
| G: edge masking + RWR | 2026-Q4 | mutation-conditioned dynamic graph | n.a. | n.a. | n.a. | edge masking hurts; RWR alone +0.004; mutation topology not bottleneck (D-076) |
| H: cross-attention | 2026-Q4 | cross-attention drug-cell fusion | 0.55 | n.a. | n.a. | collapsed; aggressive aggregation kills signal (D-078) |
| I: production (GAT+FiLM+module+RWR) | 2026-Q4 | headline architecture | 0.7723 | 0.7655 | 0.7597 | multi-seed mean (42/7/123); seed-std 0.0006/0.0006/0.0058 (D-082) |
| I + HGBR α-ensemble | 2026-Q4 | per-fold val carve-out fits scalar $\alpha$ | 0.7796 | 0.7717 | 0.7629 | $\alpha \in [0.58, 0.61]$ on hard splits; both models contribute (D-090, D-092) |

**Table S6:**
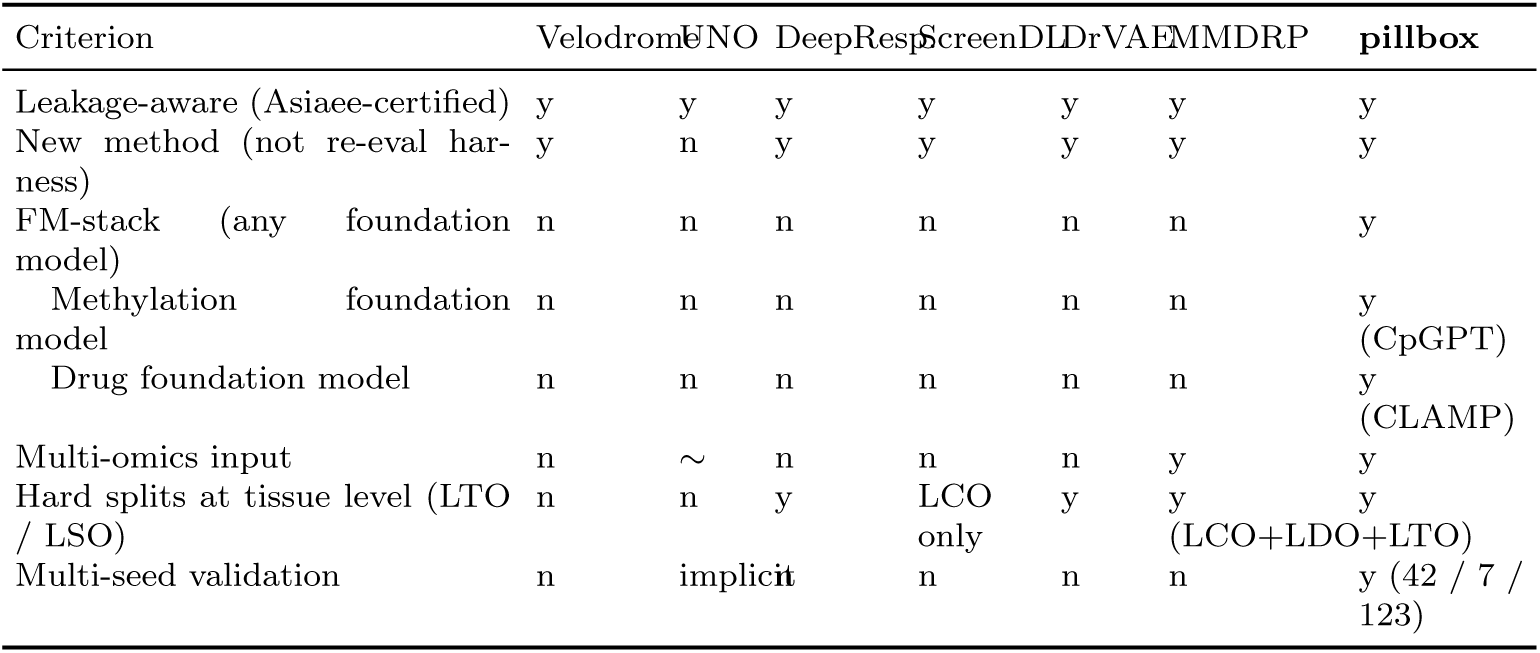
Asiaee 2026 leakage-clean roster comparison. Criteria are rows; methods are columns. “y” = satisfies criterion, “n” = does not, “~” = partial. LTO = leave-tissue-out; LSO = leave-site-out; LCO = leave-cell-out; LDO = leave-drug-out.

### S7. Full per-seed × per-fold results matrix (Supplementary Table S7)

Per-seed per-fold test R^2^ / RMSE / Pearson correlation for five Arch I variants (the baseline, the per-fold-PCA-clean ablation, the two sparse-bio ablations, and the *α*-ensemble with HGBR) across the three split types. 206 rows total. The across-seed standard deviation is approximately an order of magnitude smaller than the within-seed across-fold standard deviation at the architecture-stable convergence point (for example, on random 5-fold, Arch I shows seed-std 0.0006 and fold-std around 0.011), providing the basis for our claim that the headline numbers are not artifacts of the seed-42 draw.

Model legend for Supplementary Table S7: `I` = Arch I; `I+HGBR` = Arch I + HGBR *α*-ensemble (headline architecture); `I/PCAcln` = Arch I with the M2-clean per-fold-PCA preprocessing (Section 3); `I/bio-mut` = Arch I + sparse-bio mutations channel; `I/bio-all` = Arch I + sparse-bio all-combined channel. The last two are the negative-result curated-bio-prior ablations. `n_test` is the number of (cell, drug) pairs in the held-out test fold; “n.a.” for the `I/PCAcln` rows, which were re-trained against the original test split.

**Table S7:**
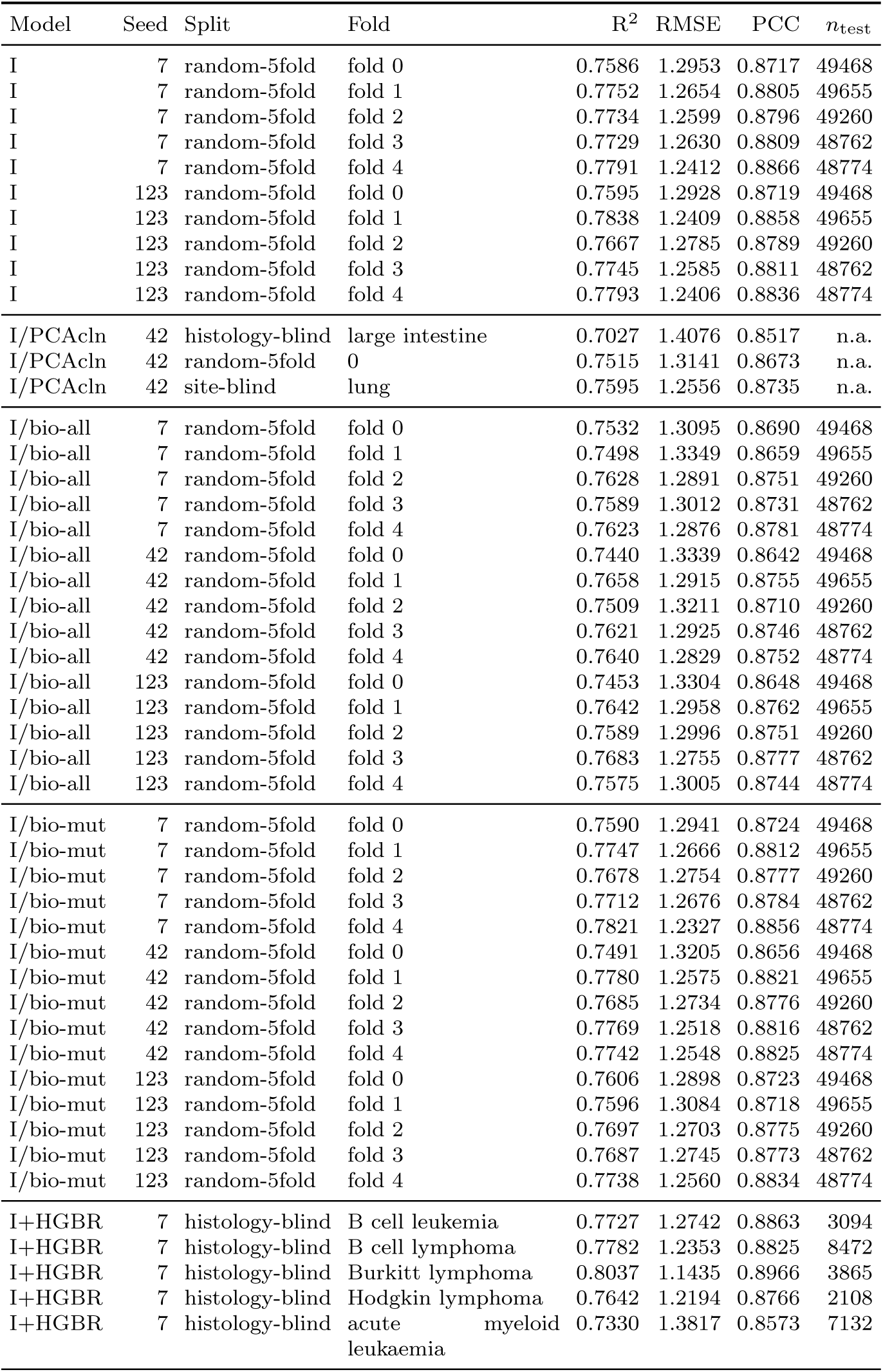
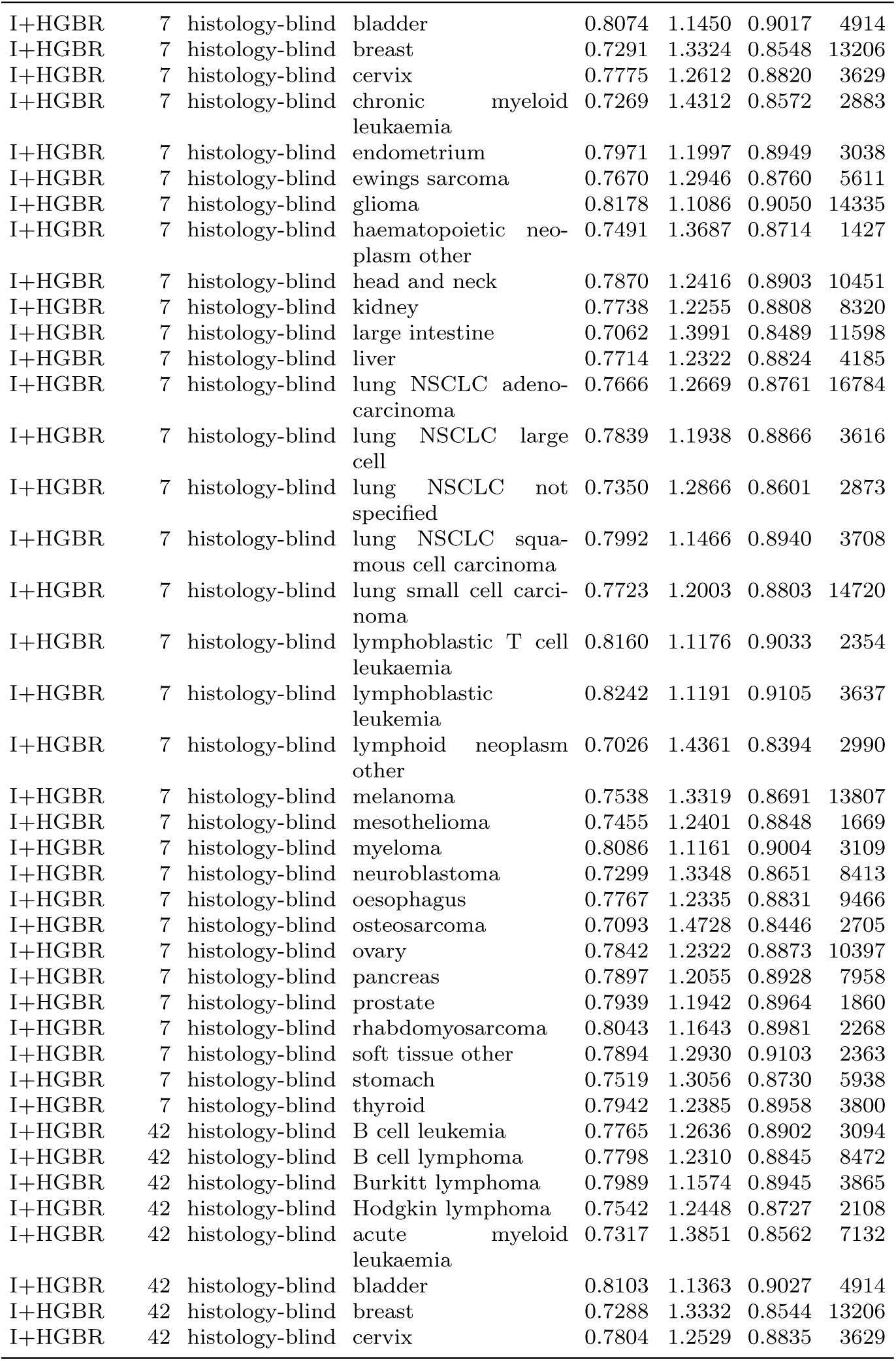
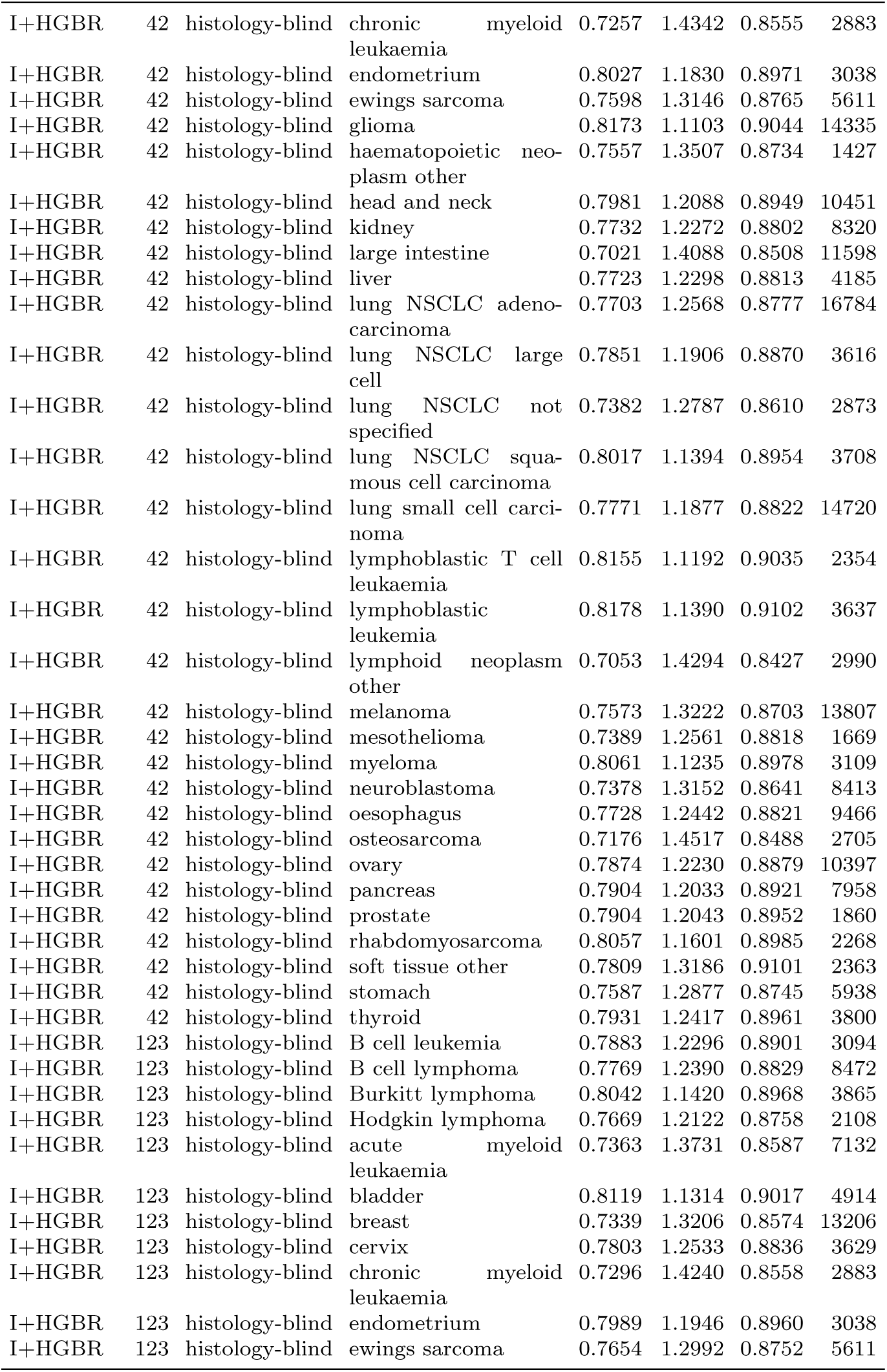
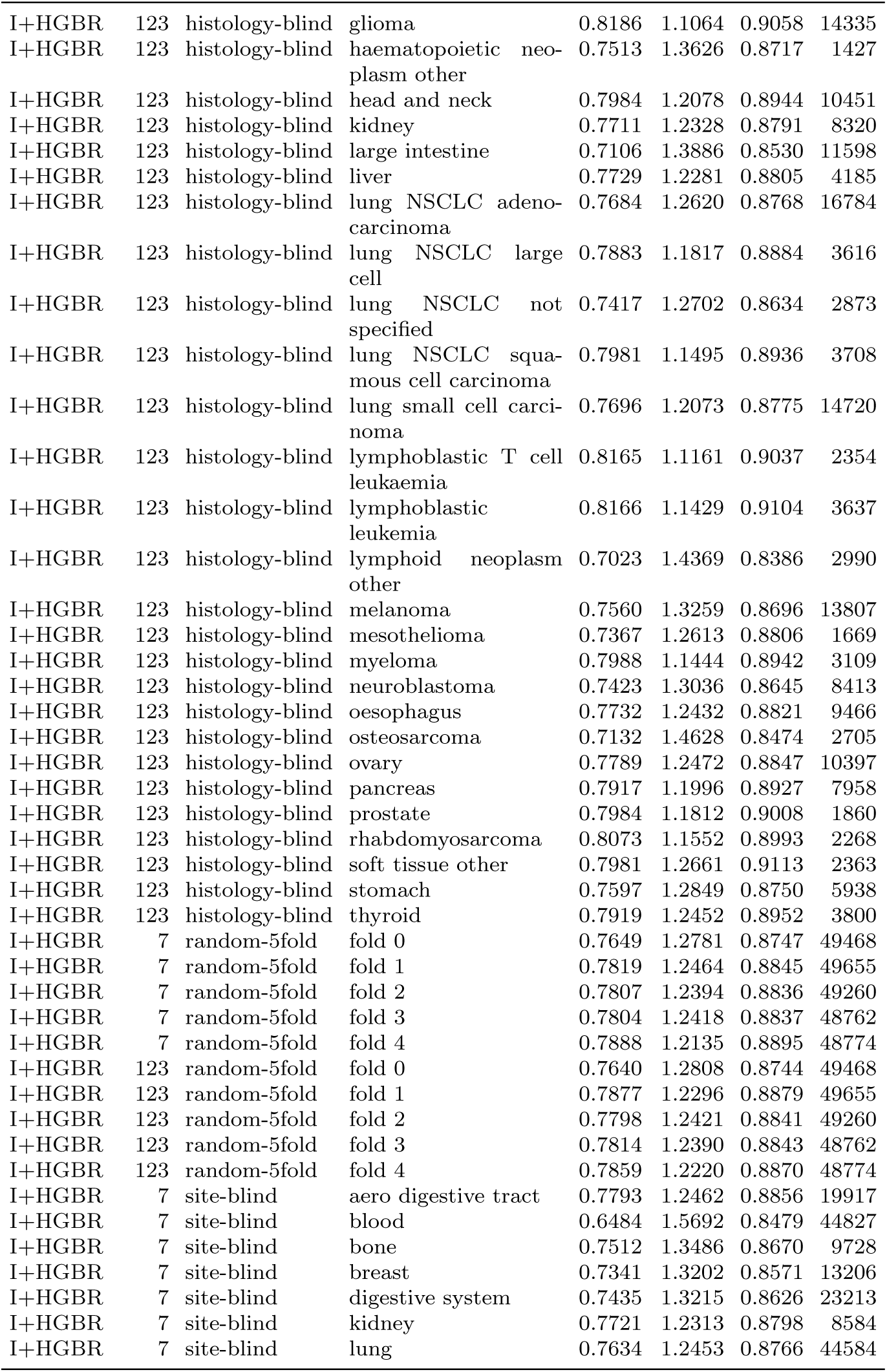
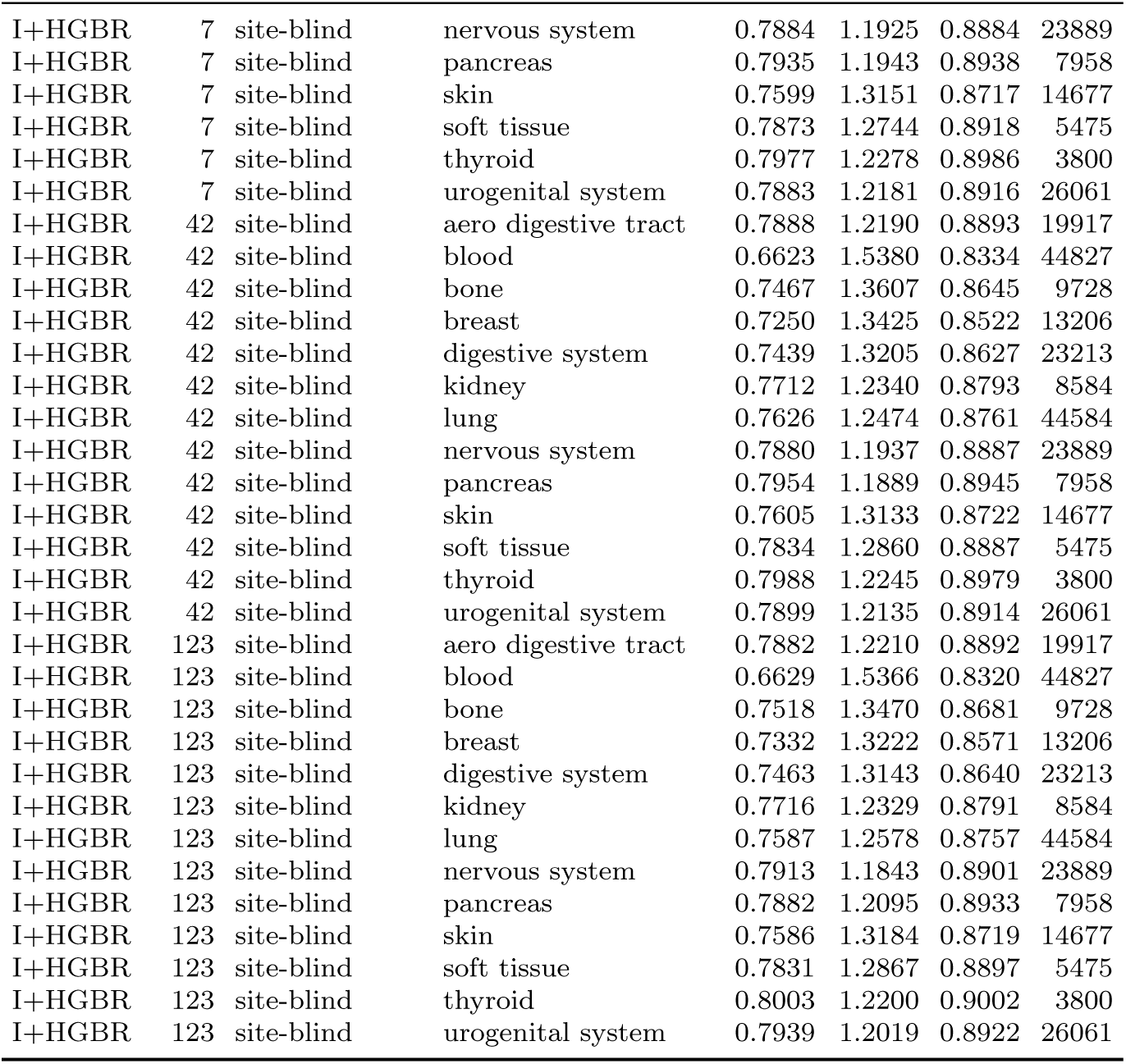
Full per-seed-per-fold results matrix (206 rows). See preceding paragraph for the model legend.

### S8. TransCDR public-code M2 leakage

The TransCDR paper [30] reports random-split Pearson 0.94 and cold-cell Pearson 0.86 on GDSC. Inspection of the public training pipeline at github.com/XiaoqiongXia/TransCDR shows that `Step1_Data_split.py` reads prez-scored expression and methylation matrices (`RNA_n18451_1018_zscore.csv` and `mrna_n20617_1028_zscore.csv`, z-scored across all 1,018 / 1,028 cell lines in the source files) before applying `KFold` cross-validation. This is Asiaee mode M2 (preprocessing fit on full data before the train/test split) and is a confirmed leakage vector in published code that we did not test against in our TransCDR-like architecture-only comparator (Section 3). Our comparator uses our own preprocessing pipeline (per-fold scaler, frozen FM embeddings); the 5–8 R^2^ gap we report against TransCDR-like is thus a reflection of fusion-architecture differences on clean inputs.

### S9. CpGPT pretraining corpus disjoint from evaluation cohort

The CpGPT foundation model [17] is pretrained on the “CpGCorpus,” approximately 150,000 methylation samples from 2,042 GEO studies, none individually enumerated in the preprint. To establish that our evaluation cohort GSE68379 was not in this corpus, we contacted the lead CpGPT author by email in April 2026; the author confirmed by reply that GSE68379 was not included in the CpGCorpus pretraining set. The CLAMP drug foundation model [18] was pretrained on independent ligand-language paired corpora (PubChem assay descriptions and similar) and has no possible overlap with drug-response label data.

### S10. CpGPT linear probing on tissue, histology, and driver mutations

To characterize what CpGPT embeddings linearly encode, we ran 5-fold cross-validation logistic regression on the frozen 512-d CpGPT cell embeddings for three classes of target: primary tissue site (13 classes, *n* = 901), primary histology (38 classes, *n* = 854), and binary driver-mutation status for 17 canonical drivers (TP53, KRAS, BRAF, PIK3CA, PTEN, EGFR, APC, RB1, CDKN2A, MYC, BRCA1, BRCA2, NRAS, STK11, NF1, ATM, CDH1).

Primary site is decodable at accuracy 0.7025 (chance 0.0769; 9.13× above chance, macro-F1 0.5945). Primary histology is decodable at accuracy 0.5539 (chance 0.0263; 21.05× above chance, macro-F1 0.3989). Among driver mutations, only TP53 shows linearly decodable signal beyond the majority baseline (accuracy 0.7004 vs majority 0.6759, macro-F1 0.7868); all 16 other drivers have F1 between 0.02 and 0.44 with accuracy within 1% of the majority baseline. This is consistent with [36]’s characterization of TP53 as a pan-cancer master regulator whose loss-of-function is reflected in downstream methylation patterns, while other driver-mutation states have effects that are less linearly decodable from cell-line methylation alone.

These probing results support the lineage-saturation interpretation of Sections 3 and 4: CpGPT cell embeddings linearly encode tissue identity densely and only one driver mutation (TP53) explicitly; the rest of the predictive subspace operates through tissue-correlated channels that are shared between CpGPT, the curated bio priors, and the rest of the foundation-model stack. Probing script: `probe_cpgpt_representations.py`; raw output: `results/cpgpt_probing/probing_results.json`.

### S11. Data and code availability

All code, foundation-model embeddings, deterministic fold definitions, per-fold prediction arrays, ensemble outputs, and intermediate result files referenced in this paper are deposited at github.com/just5034/pillbox_research_data. The repository includes the curated training tables (methylation, gene expression, and IC_50_ in joined CSV form), the foundation-model embeddings (CpGPT 512-d cell, CLAMP 768-d drug, expression PCA-128 with gene loadings), per-fold prediction arrays from Arch I and from the HGBR baseline across all three splits and all multi-seed runs, every result JSON cited in the text, all five final figures, and the seven supplementary tables. The deterministic split JSON files are at `data/training/splits/` in the same repository.

Raw methylation data is from GEO accession GSE68379. Raw GDSC1 IC_50_ values are at cancerrxgene.org. Raw DepMap 24Q4 binary mutation calls are at depmap.org. The PRISM Repurposing Secondary Screen data is at depmap.org/portal under accession PRISM Repurposing Public 19Q4 secondary screen.

Trained model weights for Arch I (per-seed, per-fold PyTorch checkpoints, approximately 1.5 GB across all splits and seeds) are available from the corresponding author on request. They are not required to reproduce any number in the paper from the deposited prediction arrays. If one wants to run new inference with the trained weights themselves, the data repository contains the training scripts and a deterministic seed protocol that will regenerate equivalent weights in roughly 80 A100 GPU-hours.

The CpGPT methylation foundation model is at github.com/lucascamillomd/CpGPT. The CLAMP drug foundation model is at github.com/ml-jku/clamp. Seed control covers `numpy, torch`, Python `random`, and CUDA. Reported numbers are reproducible within approximately 10^−4^ R^2^ tolerance due to mixed-precision bf16 autocast.

